## Supplemental Information for "Structural and functional characterization of the NF-κB-targeting toxin AIP56 from *Photobacterium damselae* subsp. *piscicida* reveals a novel mechanism for membrane interaction and translocation"

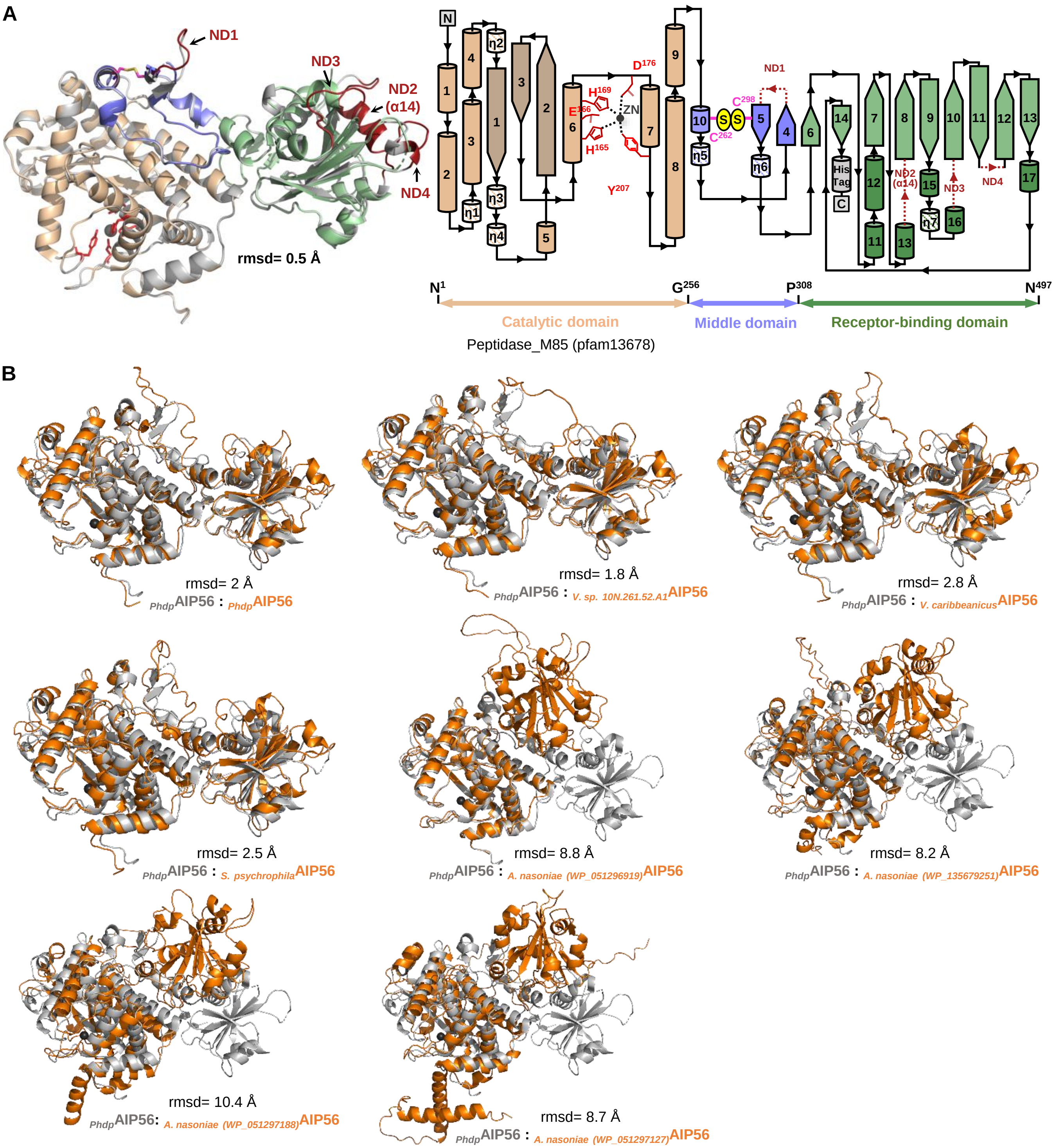


**Figure S1**: **(A)** Left: Cartoon representation of superposed AIP56 crystal structure (colored as Fig. 1A) and AIP56 model generated with Modeller program and AlphaFold2-Advanced (grey). The added regions (ND1-4) that were absent in AIP56 crystal structure are highlighted in dark red. Right: Topological representation of AIP56 with domains colored as in Fig. 1A. The regular secondary-structure elements are depicted and labeled. **(B)** Superposition of AIP56 crystal structure (grey) and AIP56 and AIP56 homologue models (see Table S2 for accession numbers) generated by AlphaFold2_Advanced (orange). Due to the high number of AIP56-like proteins from *Vibrio* species/strains, only the models with the lowest and highest rmsd are shown.

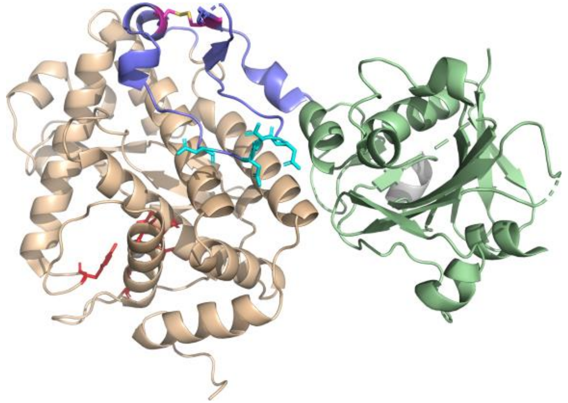

Similar to NleC

```
S. enterica NleC (WP_000704096.1)            (…) HEDNEEAFFERLDVISEGYEASADFTEYPV-MSDMVKELNKPH--DF------------PGLVIND----NT------MDADPDQIQLY-HGQPYIFTFVDKHNQR---  330 (…)
P. damselae AIP56 (WP_012954632.1)           (…) HGSELGFLFTRLATIAKGKKASPDFGTLTSFCSEGISSF--PKYPDHDD-DFNGGGAFFLPSASADSSVECTFDVLNRIEPVDDSIKFE-GGNLLIKNDFKNLNLRVAQ  335 (…)
V. splendidus (WP_032554400.1)               (…) HPSEASALLQRISIIARGSKASTSFSLLTSFCSSMNVDL--PDRPNFDDDDFSMGAAFFTGASASN-SGQCSFDVKDRAKPISDSIHFE-GGQFLIKRDLINLNLSVAK  357 (…)
V. sp. HI00D65 (WP_063524616.1)              (…) HPSEASALLQRISIIARGSKASTSFSLLTSFCSSMNVDL--PDRPNFDDDDFSMGAAFFTGASASN-SGQCSFDVKDRAKPISDSIHFE-GGQSLIKRDLINLNLSVAK  357 (…)
V. chagasii (WP_137408435.1)                 (…) HPSEASALLQRISIIARGSKASTSFSLLTSFCSSMNVDL--PDRPNFDDDDFSMGAAFFKGASASN-SDQCSFDVKDRAKPISDSIHFE-GGQFLIKRDLINLNLSVAK  357 (…)
V. lentus (WP_102413802.1)                   (…) HPSEASALLQRISIIARGSKASTSFSLLTSFCSSMNVDL--PDRPNFDDDDFSMGAAFFTGASASN-SGQCSFDVKDRAKPISDSIHFE-GGQFLIKRDLINLNLSVAK  357 (…)
V. (WP_017104811.1)                          (…) HPSEASALLQRISIIARGSKASTSFSLLTSFCSSMNVDL--PDRPNFDDDDFSMGAAFFTGASASN-SGQCSFDVKDRAKPISDSIHFE-GGQFLIKRDLINLNLSVAK  357 (…)
V. sp. 10N.261.52.A1 (WP_102424773.1)        (…) HPSEASALLQRISIIARGSKASTSFSLLTSFCSSMNVDL--PDRPNFDDDDFSMGAAFFTGASASN-SGQCSFDVKDRAKPISDSIHFE-GGQFLIKRDLINLNLSVAK  357 (…)
V. sp. 2521-89 (WP_089070319.1)              (…) HPAEAKALLSRLATIADGSNTSASFSPLTSFCSSGISEL--PKLPDFDGDDFSMGAAFFTGATASN-LGKCSLDADDRVNPVSNSITFE-GGQVLIKRDLKNLNLLVAK  357 (…)
V. sp. 2015V-1076 (WP_113597563.1)           (…) HPAEAKALLSRLATIADGSKTSASFSPLTSFCSSGISEL--PKLPDFDDDDFSMGAAFFTGATASN-LGKCSLDADDRVNPVSNSITFE-GGQVLIQRDLKNLNLLVAK  357 (…)
V. sp. 2017V-1085 (WP_113602841.1)           (…) HPAEAKALLSRLATIADGSKTSASFSPLTSFCSSGISEL--PKLPDFDDDDFSMGAAFFTGATASN-LGKCSLDADDRVNPVSNSITFE-GGQVLIQRDLKNLNLLVAK  350 (…)
V. caribbeanicus (WP_139056856.1)            (…) HPSETVALLERIATISRGLKASASFSLLTSFCSSGIIEL--PKLPDFDDDDFSMGAAFFTGASASN-LGECSLDAAGRVEPVSSSITFE-GGQLLIQRDLKNLNLLVAK  356 (…)
V. jasicida (WP_104037599.1)                 (…) HPSEAEALLNRLATISRGSKASARFSLLTSFCSSGIIDL--PKLPDFDDDDFSMGPVFFTGASASN-LGECSLDVTGRVEPVNPSITFE-GGQLLIQRDLKKLNLLVAK  357 (…)
V. sp. AND4 (WP_009841419.1)                 (…) HPYEAEALLNRLATISSGLKASPRFSLLTSFCSSGIIDL--PKLPDFDDDDFSMGAAFFTGASASS-SGSCSLDAAGRVDPVSSSITFE-GGQLLIQRDLKNLNLLVAE  357 (…)
S. psychrophila (WP_077754668.1)             (…) HPFEATELMGRMSTISAGLQASTRFDLLTSFCSSNKIDW--PPLPDFDDDDFSMGSAFFTGASASNLVDQCSIDVTKRVDTVSEAIEFE-GGQLLIQRDFKNLNLLVAK  357 (…)
V. azureus (WP_021710670.1)                  (…) HPAEASELMSRLATIASGLRASNQFTLLTSYCSSSQIDL--PPLPDFDDDHFNMGAAFFTGASASQSVGRCSIDNALRVQPVTEDIRFA-AGYPLIKRDFKNLNLLVAK  357 (…)
V. (WP_039981518.1)                          (…) HPAAASELMSRLATIASGLRASNQFTLLTSYCSSSQIDL--PPLPDFDDDHFNMGAAFFTGASASQSVGRCSIDNALRVQSVTEDIRFA-AGYPLIKRDFKNLNLLVAK  357 (…)
A. nasoniae (WP_051296919.1)                 (…) HPRRENELLERLISISENVGASSDFSSLSESSNDSSVSTAACRFEHGDSDDDFTGSTFFRGAKAANYLCH-----------KKVKIIFE-NDLSLEYWHLKKYNLIIAE  322 (…)
A. nasoniae (WP_135679251.1)                 (…) HHQRGYELLERLCDINHLLTSSPNFAG-IDTLS---DECLIPTHSSDEDTPAHWGAFLLSGAHATH----QRKDV---FATKYKNIVFNKEELPLTQWHLEKYGLVIAE  347 (…)
A. nasoniae (WP_051297188.1)                 (…) HHEHPTEVINRLFAINFFVENRPEDLNNHPPQY---FRYLLPDHLFCDDNDVQLGSAFFMGASASH----CKNNL---FAAKYENIIFS-ENLPLTQFHLEKYNLVIAA  342 (…)
A. nasoniae (WP_051297127.1)                 (…) SENL-SERVNRLLEIGDRMVAEIAAQ---PYAHEPNAEL--LSICPFDDDQEHLRFPFFSGAHGISVA-KS------FFCINHSDIIFN-DGLKLSNLHLIKYDLAAAT  403 (…)
APSE-2 Protein D (WP_015874047.1)            (…) --MEAEIGMKK---KSLNLKSIIVISVLLLLFS---------NHYTYAEKKYDFTKYCFSG------------------RP-VNHVIFN-DNTPLYEWELKALNLTIVE  75 (…)
```

Similar to APSE2 Protein D

**Figure S2**: Top: AIP56 three-dimentional structure with the conserved aspartates shown as cyan sticks. Bottom: Multiple sequence alignment (Clustal Omega: default parameters) with NleC, APSE2 Protein D, AIP56 and AIP56 homologues showing the boundaries between the catalytic, middle and receptor-binding domains. Only the region near the domain boundaries is shown. Shaded magenta: cysteine residues (in AIP56 form a disulfide bond); Shaded cyan: conserved aspartate patch within the linker peptide. Shaded grey: regions homologous to NleC and APSE2 Protein D. Note: for AIP56 the amino acid numbering considers only the mature protein (without the signal peptide), in congruence with previous publications. For the other proteins the entire proteins were considered, including the signal peptide.

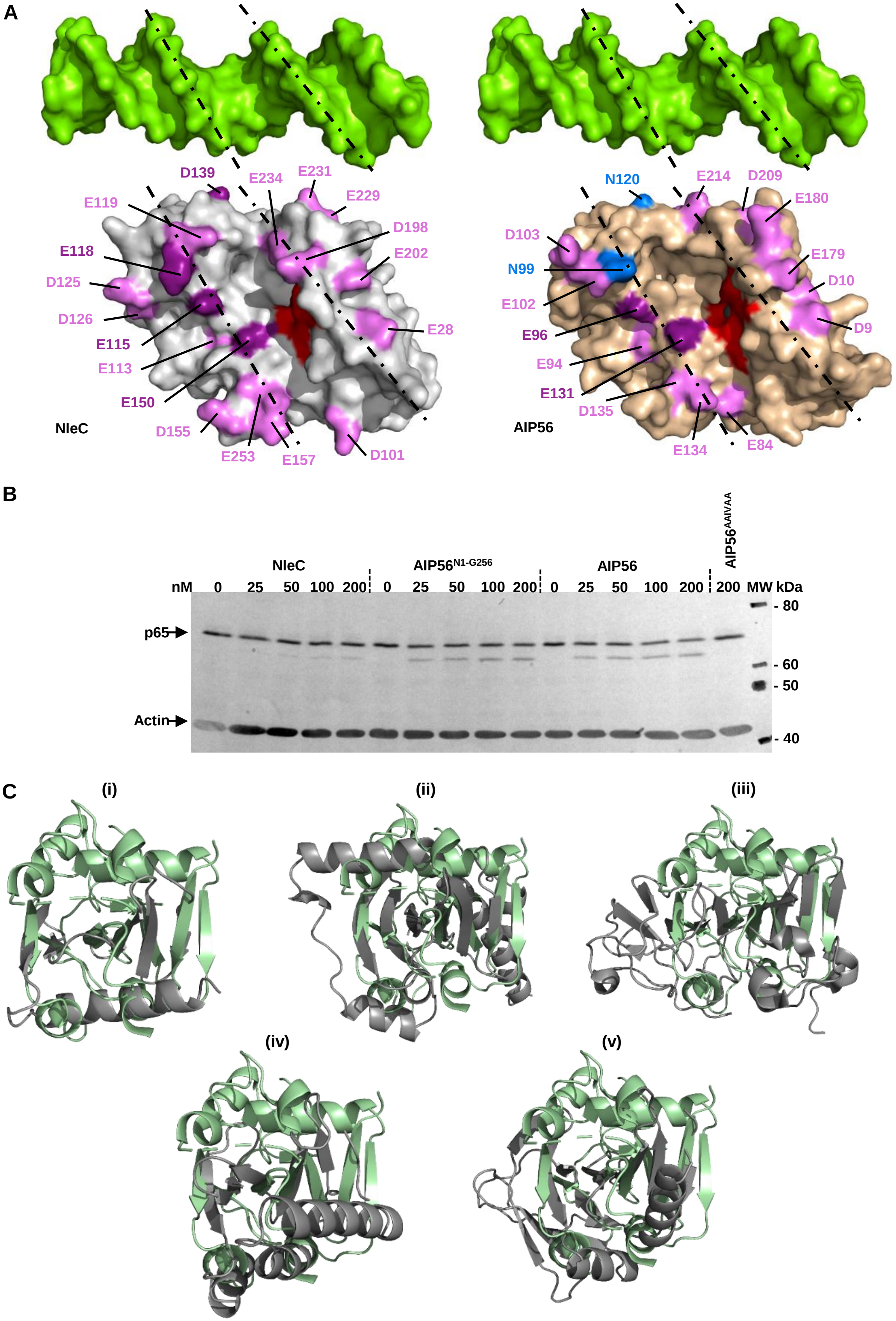
A
D139
E234
E231
E229
E119
D198
E118
E202
D125
D126
E28
E115
E113
E150
NleC
D155
E253
E157
D101
N120
E214
D209
E180
D103
E179
N99
D10
E102
D9
E96
E94
E131
D135
AIP56
E134
E84
B
AIP56AAIVAA
NleC
AIP56N1-G256
AIP56
nM 0 25 50 100 200 0 25 50 100 200 0 25 50 100 200 200 MW kDa
p65
Actin
- 80
- 60
- 50
- 40
C
(i)
(ii)
(iii)
(iv)
(v)

**Figure S3: (A) As in NleC, the active center cleft of AIP56 also mimics the major groove of DNA.** Surface representation of the DNA fragment that was complexed with NF-κB p65 (RelA) (PDB entry 1RAM), of the NleC (PDB entry 4Q3J) and of the catalytic domain of AIP56 showing the similarity between the major groove of the DNA and the cleft of the active centers of the metalloprotease. Negatively charged residues disposed along the ridge or face side of the catalytic centers are colored violet or purple. Purple colored residues are residues that have been shown to be important for the efficient proteolysis of p65 by NleC. Their counterparts in the catalytic domain of AIP56 are also highlighted in purple while the non-conserved asparagine residues are colored blue. Active site residues are colored red. **(B) AIP56 cleaves human NF-kB p65 more efficiently than NleC.** Proteolysis of NF-kB p65 by NleC from *E. coli* O157:H7 strain 4462 and AIP56 from *Photobacterium damselae* subsp. *piscicida*. HeLa lysates (20 µl) were incubated with NleC, $AIP56^{N1\text{-}G256}$ (catalytic domain), AIP56 (full length) or inactive $AIP56^{AAIVAA}$ (negative control) at the indicated concentrations for 2 h at room temperature and p65 cleavage accesses by western blotting. **(C) The twisted antiparallel β-sheet fold in AIP56 receptor-binding domain (green) is common to a number of proteins (grey),** the closest of which (as identified with PDBeFold, https://www.ebi.ac.uk/msd-srv/ssm/) are (i) the PB3 domain of PLK4 from *Drosophila melanogaster* (PDB entry 5LHZ; rmsd = 2 Å, Percentage of Sequence Identity (%Seq) = 4) (M. A. Cottee et al. 2017, *Biology Open* 6, 381-389), (ii) the hypothetical protein from *Leishmania major* homologue to human p32 protein (PDB entry 1YQF; rmsd = 3 Å, %Seq = 10), (iii) the C-terminal fragment of Zika virus nonstructural protein 1 (PDB entry 5IY3; rmsd = 3.4 Å, %Seq = 4) (H. Song, et al. 2016, *Nature Structural & Molecular Biology* 23, 456-458), (iv) the integron cassette protein VCH_CASS14 from *Vibrio cholerae* (PDB entry 3IMO; rmsd = 4 Å, %Seq = 7) (V. Sureshan et al., 2013, *PLOS ONE* 8, e52934) and (v) the invasion associated protein B from *Bartonella henselae* (PDB entry 3DTD; rmsd = 4.3 Å, %Seq = 8).

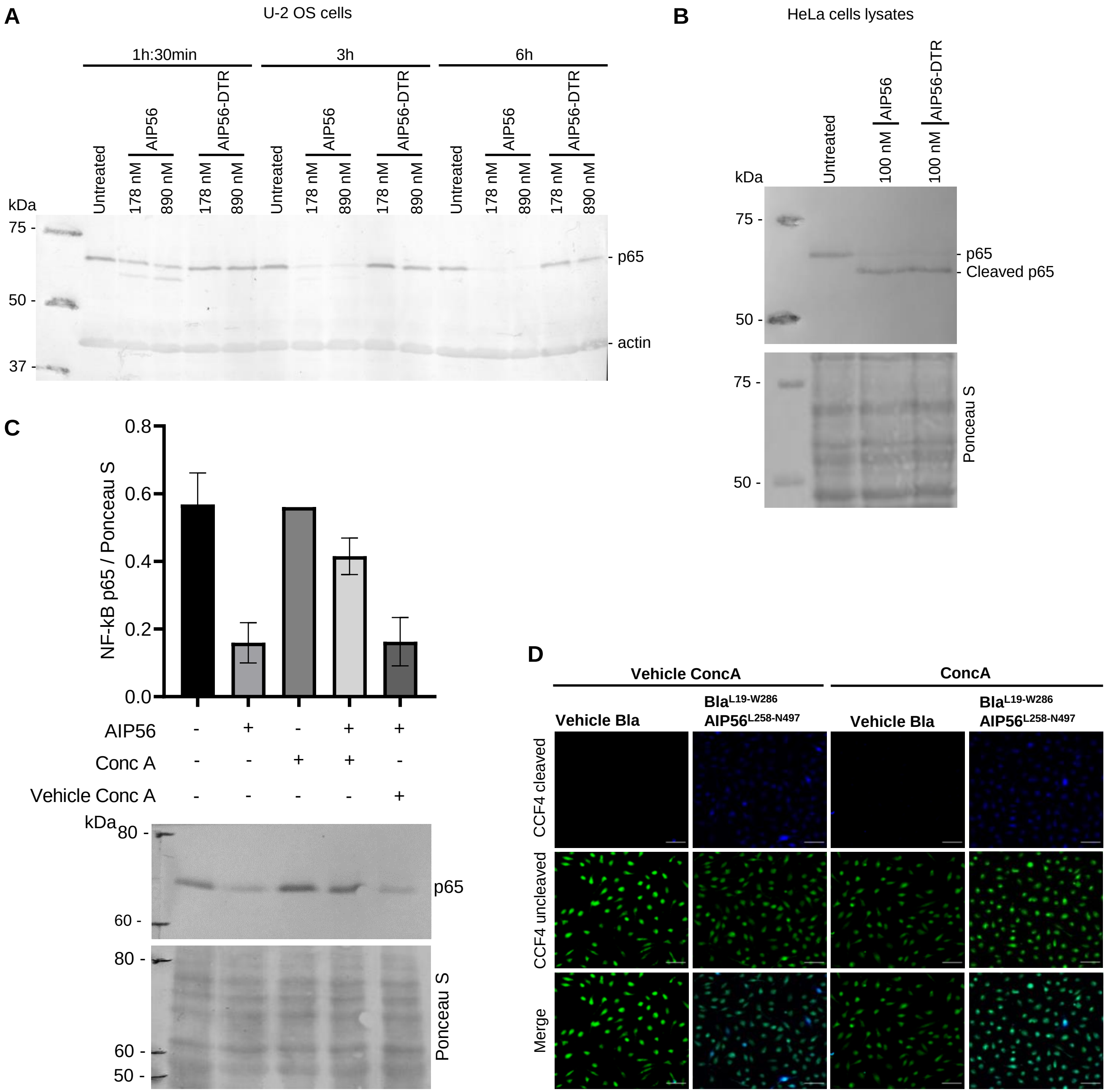
A
U-2 OS cells
1h:30min
3h
6h
Untreated
178 nM
890 nM
AIP56
AIP56-DTR
kDa
75 -
50 -
37 -
- p65
- actin
B
HeLa cells lysates
kDa
Untreated
100 nM
AIP56
AIP56-DTR
75 -
50 -
- p65
- Cleaved p65
Ponceau S
C
NF-kB p65 / Ponceau S
0.0
0.2
0.4
0.6
0.8
AIP56
Conc A
Vehicle Conc A
kDa
80 -
60 -
50 -
p65
Ponceau S
D
Vehicle ConcA
ConcA
Vehicle Bla
BlaL19-W286 AIP56L258-N497
CCF4 cleaved
CCF4 uncleaved
Merge

**Figure S4: Pore formation requires both the middle and receptor-binding domains. (A)** AIP56$^{N1-E307}$DTR$^{S406-S560}$ (AIP56-DTR) didn't cleave p65 in U-2 OS cells. Cells were seeded at a density of 2 × $10^5$ cells per well in flat-bottom 24-well plates (Thermo Scientific, 142475) and allowed to attach and grow for 24 h at 37 ºC in a humidified chamber (5% $CO_2$) in DMEM containing 10% FBS, 4 mM L-Glutamine and 1% P/S. Then, the cells were washed twice with PBS and loaded with the indicated protein in 250 µl of DMEM for the indicated time. After incubation, cells were washed twice with PBS and collected by resuspension in SDS-PAGE sample buffer. Protein samples were boiled for 5 min before SDS-PAGE and transferred onto nitrocellulose membranes for western blotting as described in section 4.13. Cleavage of p65 and loading control of actin (anti-human actin rabbit polyclonal antibody H-196, Santa Cruz Biotechnology, #sc-7210) were revealed by chromogenic detection. Blot shown is representative of two independent experiments. **(B)** AIP56$^{N1-E307}$DTR$^{S406-S560}$ (AIP56-DTR) is catalytically active. HeLa cells lysates were obtained by resuspending 2 × $10^5$ cells on ice in 10 mM Tris pH 8.0, 150 mM NaCl, 0.5% (v/v) Triton-X100, 10% (v/v) glycerol. 20 µl of lysate was incubated with 100 nM of the indicated proteins for 2 h at RT. After incubation, samples were prepared for western blotting as described in section 4.13. Cleavage of p65 was revealed by chromogenic detection and protein loading was controlled by Ponceau S staining. **(C)** Control of ConcA activity by confirming its inhibitory effect on NF-kB p65 cleavage upon AIP56 intoxication of mBMDM. As previously described (*Infect. Immun.* 2014, 82(12):5270. doi: 10.1128/IAI.02623-14), AIP56 requires endosomal acidification to reach the cytosol and cleave NF-kB p65. A representative blot of two independent experiments is shown. Loading correction was achieved by dividing the density of p65 by the respective density of the Ponceau S staining. Values are mean ± SD. **(F)** Representative images used for the quantification shown in Fig. 2C. Scale bar = 50 µm.

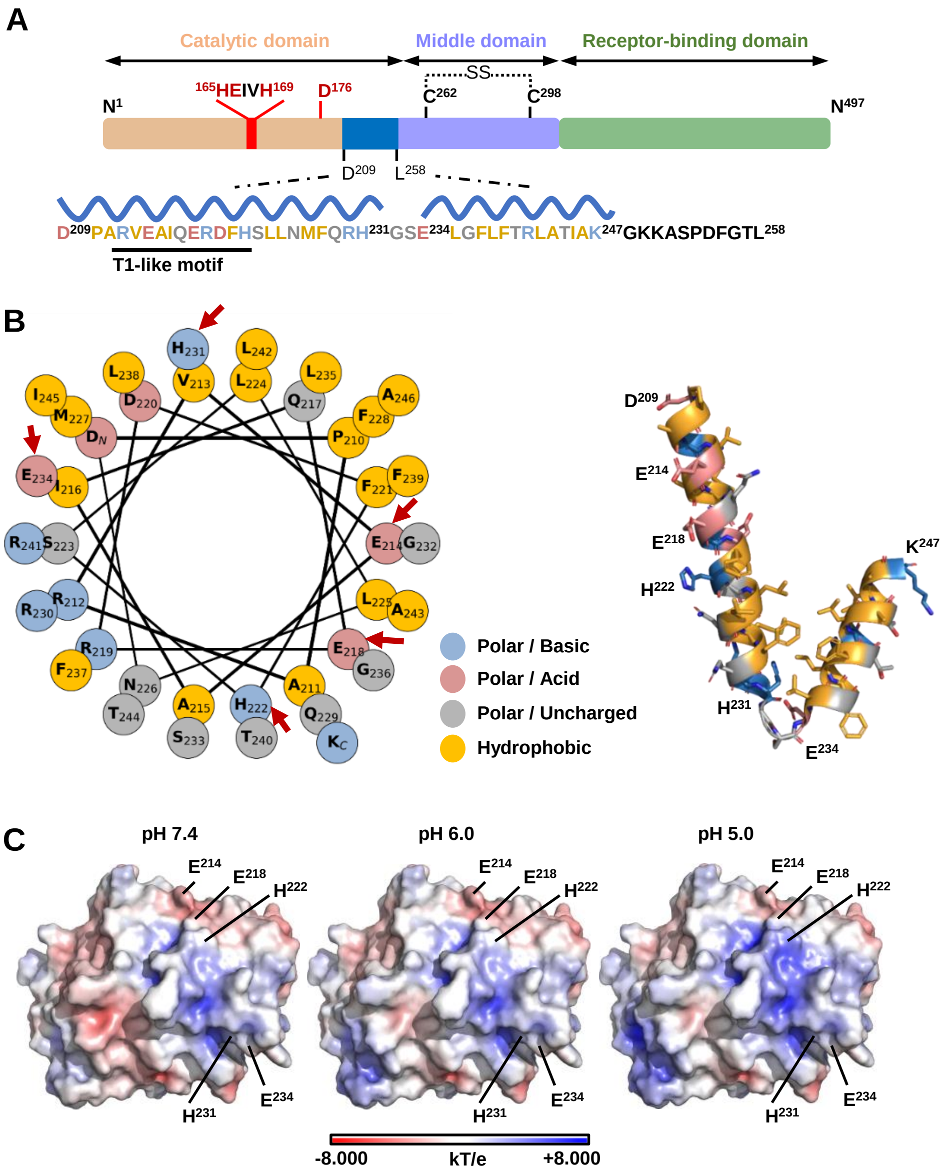


**Figure S5. Schematic representation of AIP56 structural domains, hydrophobicity analyzes of the two helices in the D209-L258 region, and surface charge distribution analysis.** (A) Schematic linear representation of AIP56 structural domains. The domains are colored as in Fig. 1A; D209-L258 region (marine blue) including two helices (D209 to H231 and E234 to K247) and a conserved T1-like motif (underlined); zinc metalloprotease catalytic center (red). **(B)** Left: Galaxy helical wheel projection (https://cpt.tamu.edu/galaxy-pub) of the two hydrophobic helices forming the D209-K247 hairpin. Residues potentially involved in pH-sensing are highlighted with red arrows. Right: Cartoon representation of the D209-K247 hairpin. Their amphipathic nature is shown, as hydrophobic residues (orange sticks) are concentrated on one side whereas basic residues (blue sticks) and hydroxylated polar residues (grey sticks) form the polar face of the helices. **(C)** Surface charge distribution (blue, positive; red, negative; white, uncharged) of the catalytic domain of AIP56 calculated at pH 7.4, 6.0 and 5.0. The negative surface charge became more neutral/positive with decreasing pH, except for residue $E^{214}$.

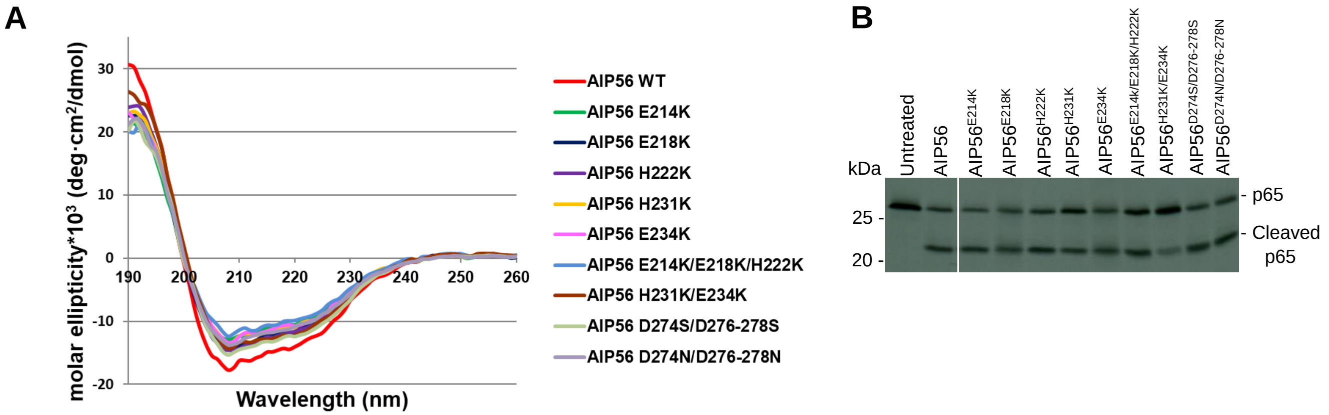


| | WT | E214K | E218K | H222K | H231K | E234K | E214K/E218K/ H222K | H231K/ E234K | D274S/ D276- 278S | D274N/ D276- 278N |
|---|---|---|---|---|---|---|---|---|---|---|
| Helix (%) | 73 | 77 | 74 | 73 | 72 | 74 | 72 | 76 | 77 | 76 |
| Strand (%) | 12 | 10 | 10 | 10 | 10 | 9 | 11 | 10 | 10 | 10 |

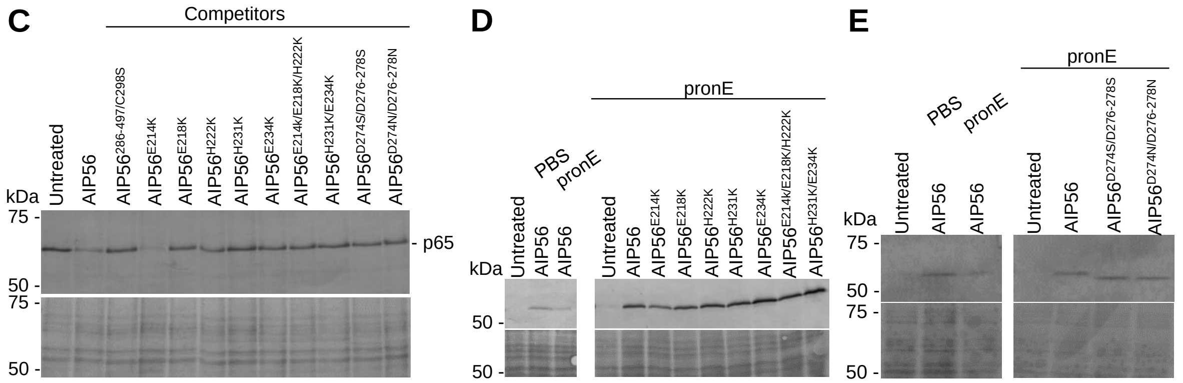


**Figure S6: All AIP56 variants are structurally stable, catalytically active and endocytosed into mBMDM. (A)** Circular Dichroism (CD) spectroscopy showing that all AIP56 variants are structurally similar to the wild type toxin. The table shows the percentage of α-helix and β-strand calculated with DichroWeb (Miles *et al. Protein Science* 2021, **31**(1):37-46, doi:10.1002/pro.4153; Whitmore and Wallace *Biopolymers* 2008, **89**:392-400, doi:10.1002/bip.20853; Whitmore and Wallace *Nucleic Acids Res* 2004, **32**:W668–73, doi:10.1093/nar/gkh371. **(B)** All variants cleave cell-free p65 Rel homology domain. Autoradiography of $^{35}$S-labeled sea bass p65 Rel homology domain incubated for 2 h at 22 ºC with 10 nM of the indicated proteins (for additional details see Silva *et al. PLoS Pathog.* 2013;9(2):e1003128. doi: 10.1371/journal.ppat.1003128). **(C)** With the exception of $AIP56^{E214K}$, which cleaves p65 because it translocates to the cytosol (see Fig. 3B), all other variants compete with AIP56 for cell internalization. $AIP56^{286-497/C298S}$ was used as positive control. mBMDM were pre-incubated for 15 min on ice with 35 µM of each competitor, followed by incubation for further 30 min on ice with 87.5 nM of AIP56 in the presence of each competitor, followed by 10 min at 37 ºC. Unbound proteins were removed and cells were incubated at 37 ºC for 2 h. NF-kB p65 cleavage was assessed by western blotting. The results shown are representative of at least three independent experiments. **(D)** and **(E)** All variants are endocytosed by mBMDM. Cells were incubated with V5 tagged AIP56 or variants for 30 min on ice plus 10 min at 37 ºC, washed with PBS, treated with Pronase E (pronE) to remove surface-exposed toxin and analyzed by western blotting to detect intracellular toxin (anti-V5, upper lane; chromogenic detection). Cells treated with AIP56 for 30 min on ice and treated with pronE or PBS were used to control pronE efficacy (for additional details see Pereira *et al. Infect Immun.* 2014;82(12):5270-85. doi: 10.1128/IAI.02623-14). Ponceau S staining was used to control protein loading (lower panels of C, D and E).

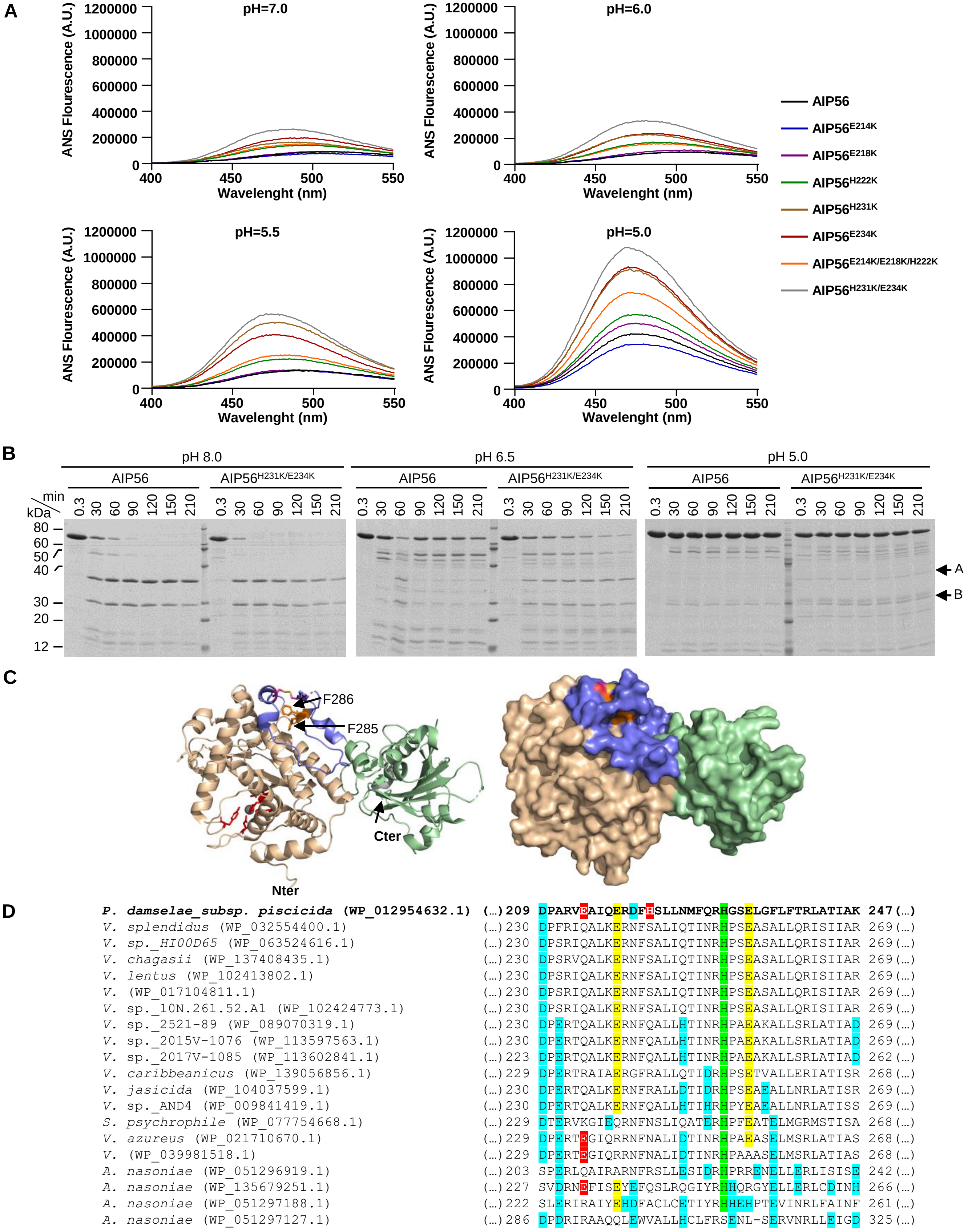
A
pH=7.0
pH=6.0
pH=5.5
pH=5.0
ANS Flourescence (A.U.)
Wavelenght (nm)
AIP56
AIP56E214K
AIP56E218K
AIP56H222K
AIP56H231K
AIP56E234K
AIP56E214K/E218K/H222K
AIP56H231K/E234K
B
pH 8.0
pH 6.5
pH 5.0
AIP56
AIP56H231K/E234K
min
kDa
0.3 30 60 90 120 150 210
80
60
50
40
30
20
12
A
B
C
F286
F285
Cter
Nter
D
P. damselae_subsp. piscicida (WP_012954632.1) (...) 209 DPARVEAIQERDFHSLLNMFQRHGSELGFLFTRLATIAK 247 (...)
V. splendidus (WP_032554400.1) (...) 230 DPFRIQALKERNFSALIQTINRHPSEASALLQRISIIAR 269 (...)
V. sp._HI00D65 (WP_063524616.1) (...) 230 DPSRIQALKERNFSALIQTINRHPSEASALLQRISIIAR 269 (...)
V. chagasii (WP_137408435.1) (...) 230 DPSRVQALKERNFSALIQTINRHPSEASALLQRISIIAR 269 (...)
V. lentus (WP_102413802.1) (...) 230 DPSRIQALKERNFSALIQTINRHPSEASALLQRISIIAR 269 (...)
V. (WP_017104811.1) (...) 230 DPSRIQALKERNFSALIQTINRHPSEASALLQRISIIAR 269 (...)
V. sp._10N.261.52.A1 (WP_102424773.1) (...) 230 DPSRIQALKERNFSALIQTINRHPSEASALLQRISIIAR 269 (...)
V. sp._2521-89 (WP_089070319.1) (...) 230 DPERTQALKERNFQALLHTINRHPAEAKALLSRLATIAD 269 (...)
V. sp._2015V-1076 (WP_113597563.1) (...) 230 DPERTQALKERNFQALLHTINRHPAEAKALLSRLATIAD 269 (...)
V. sp._2017V-1085 (WP_113602841.1) (...) 223 DPERTQALKERNFQALLHTINRHPAEAKALLSRLATIAD 262 (...)
V. caribbeanicus (WP_139056856.1) (...) 229 DPERTRAIAERGFRALLQTIDRHPSETVALLERIATISR 268 (...)
V. jasicida (WP_104037599.1) (...) 230 DPERTQALKERNFRALLDTIDRHPSEAEALLNRLATISR 269 (...)
V. sp._AND4 (WP_009841419.1) (...) 230 DPERTQALKERNFQALLHTIHRHPYEAEALLNRLATISS 269 (...)
S. psychrophile (WP_077754668.1) (...) 229 DTERVKGIEQRNFNSLIQATERHPFEATELMGRMSTISA 268 (...)
V. azureus (WP_021710670.1) (...) 229 DPERTEGIQRRNFNALIDTINRHPAEASELMSRLATIAS 268 (...)
V. (WP_039981518.1) (...) 229 DPERTEGIQRRNFNALIDTINRHPAAASELMSRLATIAS 268 (...)
A. nasoniae (WP_051296919.1) (...) 203 SPERLQAIRARNFRSLLESIDRHPRRENELLERLISISE 242 (...)
A. nasoniae (WP_135679251.1) (...) 227 SVDRNEFISEYEFQSLRQGIYRHHQRGYELLERLCDINH 266 (...)
A. nasoniae (WP_051297188.1) (...) 222 SLERIRAIYEHDFACLCETIYRHHEHPTEVINRLFAINF 261 (...)
A. nasoniae (WP_051297127.1) (...) 286 DPDRIRAAQQLEWVALLHCLFRSENL-SERVNRLLEIGD 325 (...)

**Figure S7: (A) Protonatable residues in the D209-K247 hairpin control the low-pH triggered conformational changes in AIP56.** Representative ANS measurement curves for each pH at different wavelengths. Analysis of the conformational changes at the peak ANS fluorescence is shown in Fig. 3C. **(B) Limited proteolysis of AIP56 and AIP56$^{H231K/E234K}$ by α-chymotrypsin type II at different pH.** Coomassie Blue-stained SDS-PAGE gel of 3 µg AIP56 and AIP56$^{H231K/E234K}$ following incubation with 6.25 µg mL$^{-1}$ α-chymotrypsin type II over 210 min on ice at pH 8.0 in 20 mM Tris pH 8.0 + 200 mM NaCl, at pH 6.5 in 20 mM Bis-Tris pH 6.5 + 200 mM NaCl and at pH 5.0 in 20 mM Bis-Tris pH 5.0 + 200 mM NaCl. A and B mark the bands corresponding to the catalytic and receptor-binding domains, respectively. **(C) The bond between F285 and F286 is protected from chymotrypsin cleavage.** N-terminal Edman sequencing revealed that chymotrypsin cleavage occurred between F285 and F286 (Silva et al., 2013, PLoS Pathog 9(2): e1003128; doi:10.1371/journal.ppat.1003128). **Left:** Cartoon representation of AIP56 monomer (Chain A). F285 and F286 are represented as orange sticks. **Right:** Surface representation of AIP56 structure colored as in Fig. 1A. **(D) Multiple sequence alignment (Clustal Omega: default parameters) of AIP56 hairpin region and homologous region in AIP56-like proteins.** Shaded green: AIP56 H231 and conserved histidine residues in AIP56 homologues; shaded yellow: AIP56 E218 and E234 and conserved glutamate residues in AIP56 homologues; shaded red: E214 and conserved glutamate residues in AIP56 homologues; shaded cyan: putative protonatable residues within the hairpin region not tested in this study. Note: For AIP56 the amino acid numbering considers only the mature protein (without the signal peptide), in congruence with previous publications. For the other proteins the entire ORFs were considered for amino acid numbering.

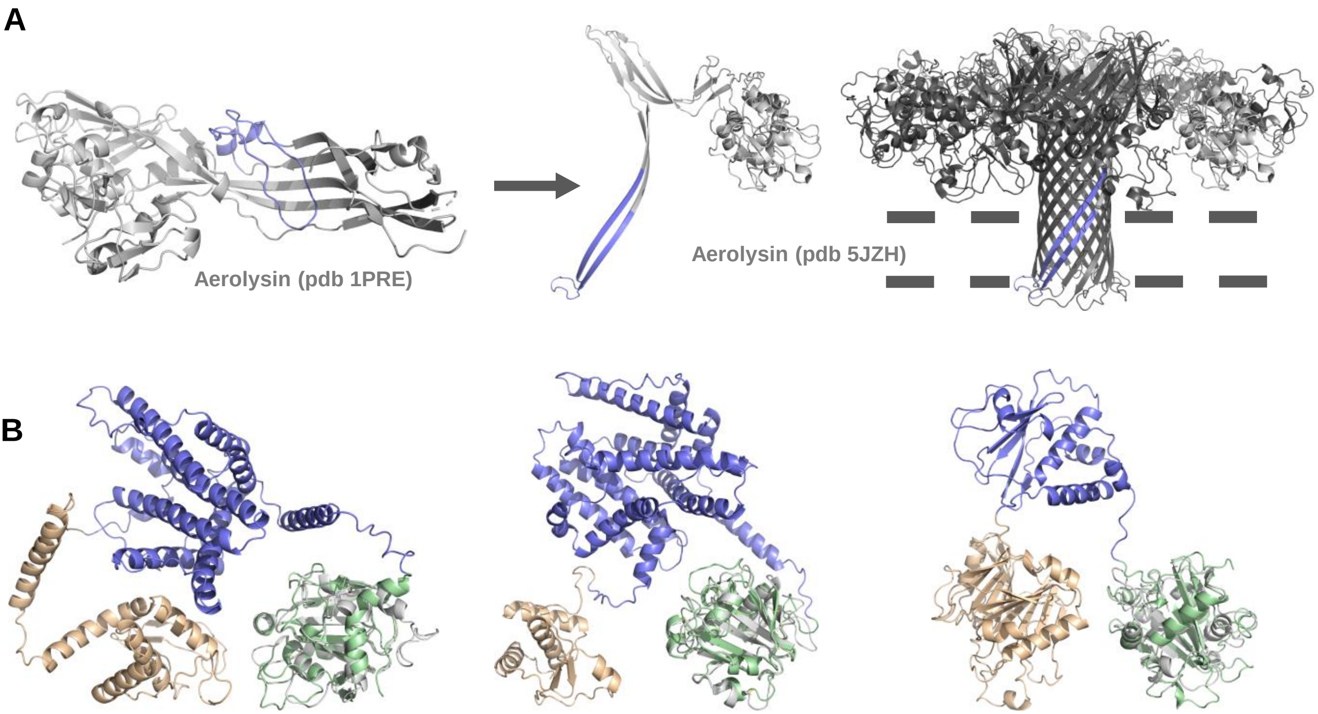


*Shewanella woodyi* AIP56-related toxin ACA88541 (colored) superposed with AIP56's receptor binding domain (grey)

*Enterovibrio norvegicus* AIP56-related toxin *WP_017014894.1* (colored) superposed with AIP56's receptor binding domain (grey)

*Drosophila bipectinata* AIP56-related toxin XP_017099943.1 (colored) superposed with AIP56's receptor binding domain (grey)

**Figure S8. (A)** AP56's middle domain is structurally similar to the region (insertion loop, prestem loop or tongue; colored slate) that in aerolysin refolds to the β-hairpin that participates in the formation of a β-barrel transmembrane channel.
**(B)** AIP56-related toxins have a middle domain with a structure characteristic of translocation domains of other short-trip single-chain AB toxins, suggesting that AIP56's receptor-binding domain and its homologous domains are not involved in pore-formation. Structures predicted by Alphfold2_Advanced and superposed with AIP56 receptor-binding domain using PyMol.

**Table S1**: Crystallography and SAXS data collection and analysis.

| Crystallography | |
|---|---|
| Data collection | |
| Space Group | $P12_11$ |
| Wavelength (Å) | 0.972 |
| Cell dimensions | |
| a,b,c (Å) | 72, 194, 92 |
| α, β, γ (º) | 90, 113, 90 |
| Resolution (Å) | 46.3 - 2.5 (2.78 - 2.5) |
| Number of observations measured | 266 172 (13 227) |
| Number of unique reflections measured | 52 821 (2 640) |
| Multiplicity | 5 (5) |
| Completeness (spherical; %) | 69.8 (14.8) |
| Completeness (ellipsoidal; %) | 93.4 (62.2) |
| I/σI | 6.7 (1.4) |
| Wilson B-factor | 48.9 |
| $R_{merge}$ | 0.163 (1.036) |
| $R_{pim}$ | 0.081 (0.509) |
| CC (1/2) (%) | 99.2 (48.8) |
| Monomers per asymmetric unit | 4 |
| Matthews coefficient ($Å^3$ $Da^{-1}$) | 2.57 |
| Solvent content (%) | 52.24 |
| Refinement | |
| $R_{work}/R_{free}$ (%) | 0.2396/0.2823 |
| Numbers of non-hydrogen atoms | 14 880 |
| macromolecules | 14 635 |
| ligands | 38 |
| waters | 207 |
| Protein residues | 1810 |
| RMSD from standard stereochemistry | |
| Bond lengths (Å) | 0.013 |
| Bond angles (º) | 1.6 |
| Ramachadran plot statistics | |
| Favored (%) | 94.4 |
| Allowed (%) | 5.03 |
| Disallowed (%) | 0.57 |
| **PDB Code** | **7ZPF** |

| SAXS | |
|---|---|
| Data collection parameters | |
| Instrument | SWING beamline (SOLEIL, France) |
| Detector | Eiger 4M Dectris |
| Beam geometry ($mm^2$) | 0.5 x 0.2 |
| Wavelenght (Å) | 1.033204 |
| q-range ($Å^{-1}$) | 0.005 - 0.6 |
| Exposure time (s) | 1 |
| SEC-SAXS column | Bio Sec 3 Agilent |
| Temperature (K) | 288 |
| Concentration range ($mg.mL^{-1}$) | 3.53 and 13.37 |
| Structural parameters | |
| $R_g$ (Å) (from P(r)) | 28.6 ± 0.2 |
| q-range ($Å^{-1}$) | 0.005-0.34 |
| $R_g$ (Å) (from Guinier plot) | 28.3 ± 0.2 |
| $qR_g$-range | 0.28-1.29 |
| $D_{max}$ (Å) | 95 ± 5 |
| Molecular mass (MM) determination (kDa) | |
| From Porod volume | 57.5 |
| From consensus Bayesian assessment | 55.6 ± 6 |
| Calculated monomeric MM from sequence | 57.25 |
| Software employed | |
| Primary data reduction | FOXTROT |
| Data processing | PRIMUS |
| Validation and averaging | CORMAP |
| Flexibility modelling | SREFLEX |
| Computation of model intensities | CRYSOL |
| 3D graphics representations | PyMOL |
| **SASBDB Code** | **SASDNW6** |

**Table S2**: Percent Identity Matrix for AIP56-like toxins from different bacterial species (created by Clustal Omega)

| | 1 | 2 | 3 | 4 | 5 | 6 | 7 | 8 | 9 | 10 | 11 | 12 | 13 | 14 | 15 | 16 | 17 | 18 | 19 | 20 |
|---|---|---|---|---|---|---|---|---|---|---|---|---|---|---|---|---|---|---|---|---|
| 1: *Phdp* WP_012954632.1 | 100.00 | 55.69 | 55.29 | 56.08 | 55.69 | 55.49 | 55.49 | 60.78 | 60.20 | 60.39 | 57.65 | 59.02 | 58.04 | 54.51 | 53.33 | 52.55 | 38.91 | 34.62 | 34.69 | 30.51 |
| 2: *Vibrio splendidus* WP_032554400.1 | | 100.00 | 96.33 | 96.53 | 97.49 | 97.49 | 97.30 | 71.24 | 71.81 | 72.60 | 70.93 | 71.43 | 70.66 | 67.12 | 62.48 | 61.51 | 41.21 | 36.25 | 37.96 | 30.40 |
| 3: *Vibrio* sp. HI00D65 WP_063524616.1 | | | 100.00 | 97.49 | 98.07 | 98.07 | 97.68 | 70.46 | 71.04 | 71.82 | 69.96 | 70.85 | 70.46 | 66.92 | 63.06 | 62.09 | 41.42 | 36.25 | 37.76 | 31.00 |
| 4: *Vibrio chagasii WP_137408435.1* | | | | 100.00 | 98.26 | 98.26 | 97.88 | 71.04 | 71.62 | 72.41 | 69.96 | 70.66 | 69.88 | 67.89 | 62.86 | 61.90 | 41.63 | 36.66 | 37.35 | 30.80 |
| 5: *Vibrio lentus WP_102413802.1* | | | | | 100.00 | 99.61 | 98.46 | 70.85 | 71.43 | 72.21 | 70.35 | 71.62 | 70.66 | 67.31 | 63.06 | 62.09 | 41.42 | 36.46 | 37.55 | 31.00 |
| 6: *Vibrio* WP_017104811.1 | | | | | | 100.00 | 98.46 | 70.85 | 71.43 | 72.21 | 70.35 | 71.24 | 70.66 | 67.50 | 63.06 | 62.09 | 41.42 | 36.46 | 37.55 | 31.00 |
| 7: *Vibrio* sp. 10N.261.52.A1 WP_102424773.1 | | | | | | | 100.00 | 71.43 | 72.01 | 72.60 | 70.74 | 71.62 | 71.04 | 66.92 | 62.86 | 61.90 | 41.21 | 36.46 | 37.96 | 30.80 |
| 8: *Vibrio* sp. 2521-89 WP_089070319.1 | | | | | | | | 100.00 | 97.10 | 97.26 | 80.04 | 81.27 | 78.57 | 67.31 | 64.60 | 63.64 | 43.51 | 39.10 | 40.20 | 31.80 |
| 9: *Vibrio* sp. 2015V-1076 WP_113597563.1 | | | | | | | | | 100.00 | 99.80 | 80.43 | 81.85 | 79.34 | 68.09 | 64.41 | 63.44 | 42.68 | 39.31 | 40.20 | 31.80 |
| 10: *Vibrio* sp. 2017V-1085 WP_113602841.1 | | | | | | | | | | 100.00 | 80.63 | 82.58 | 80.04 | 68.63 | 64.90 | 64.12 | 42.89 | 39.31 | 40.20 | 32.25 |
| 11: *Vibrio caribbeanicus* WP_139056856.1 | | | | | | | | | | | 100.00 | 82.95 | 81.20 | 67.25 | 62.72 | 61.75 | 41.96 | 39.23 | 41.14 | 32.00 |
| 12: *Vibrio jasicida* WP_104037599.1 | | | | | | | | | | | | 100.00 | 86.87 | 68.86 | 64.41 | 63.44 | 43.51 | 39.71 | 41.22 | 32.20 |
| 13: *Vibrio* sp. AND4 WP_009841419.1 | | | | | | | | | | | | | 100.00 | 68.28 | 64.22 | 63.25 | 42.68 | 39.71 | 40.41 | 32.00 |
| 14: *Shewanella psychrophila* WP_077754668.1 | | | | | | | | | | | | | | 100.00 | 70.27 | 69.50 | 42.12 | 39.15 | 39.96 | 31.75 |
| 15: *Vibrio azureus* WP_021710670.1 | | | | | | | | | | | | | | | 100.00 | 97.49 | 41.84 | 37.76 | 38.37 | 31.60 |
| 16: *Vibrio* WP_039981518.1 | | | | | | | | | | | | | | | | 100.00 | 41.84 | 37.96 | 37.76 | 32.00 |
| 17: *Arsenophonus nasoniae* WP_051296919.1 | | | | | | | | | | | | | | | | | 100.00 | 43.37 | 42.32 | 35.59 |
| 18: *Arsenophonus nasoniae* WP_135679251.1 | | | | | | | | | | | | | | | | | | 100.00 | 49.60 | 35.37 |
| 19: *Arsenophonus nasoniae* WP_051297188.1 | | | | | | | | | | | | | | | | | | | 100.00 | 38.78 |
| 20: *Arsenophonus nasoniae* WP_051297127.1 | | | | | | | | | | | | | | | | | | | | 100.00 |
